# Constitutively-active Positive Transcription Factor b (P-TEFb) prevents viral latency

**DOI:** 10.64898/2026.09.26.754700

**Authors:** Fang Huang, Kenji Nishiura, Koh Fujinaga

## Abstract

While many cellular and viral mechanisms are associated with proviral latency, the positive transcription factor b (P-TEFb), a cellular co-factor of the nuclear factor kappa B (NFκB) and viral transactiator proteins, plays a critical role in low viral transcription levels in resting infected cells. P-TEFb, comprised of CDK9 and Cyclin T1 (CycT1), is absent in quiescent CD4+T cells that represent the bulk of the viral latent reservoir. In these cells, the CycT1 subunit is dephosphorylated and dissociated from CDK9, leading to its rapid proteasomal degradation. Based on the knowledge of CycT1 phosphorylation and proteolysis, we designed a mutant CycT1 protein [constitutively active (CA)-CycT1] that is resistant to protein degradation and is stably expressed in resting CD4+ T cells. Interestingly, the expression of CA-CycT1 significantly delayed the establishment of viral latency in CD4+ T cells. This is the first example of expressing a high level of CycT1 proteins in resting CD4+ T cells, which modulates viral latency and T cell functions. This finding presents a significant therapeutic potential for the development of effective therapies for latent viral infections.

## Introduction

Although anti-retroviral therapy (ART) can reduce the replication of the human immunodeficiency type-1 (HIV) to undetectable levels, latent reservoirs remain scattered throughout the body in people living with HIV (PLWH) [1–3]. These latent reservoirs will produce fully infectious virus upon secession of the treatment, causing severe viremia leading to AIDS. Therefore, removing these latent reservoirs is a critical mandatory step to achieve a functional cure of HIV/AIDS [4, 5]. HIV latently infected cells express no detectable viral proteins, which allows these cells to escape the host immune surveillance [6]. Particularly, viral mRNA transcription is kept at a low level [7]. HIV transcription occurs from the transcription start site (TSS) located in the 5’ long-terminal repeat (LTR) of the integrated HIV provirus. In the absence of the viral Tat protein, HIV transcription is suppressed mainly at the elongation step since RNA polymerase II (RNAPII) is paused at the 5’-proximal region even it initiates transcription [8]. Tat, together with the cellular positive transcription elongation factor b (P-TEFb) releases the paused RNAPII and stimulates HIV transcription [8, 9].

P-TEFb, comprised of cyclin-dependent kinase (CDK) 9 and cyclin (Cyc) T1, T2a or T2b, is a critical transcriptional CDK/cyclin complex that stimulates the elongation step of transcription [10, 11]. P-TEFb is highly expressed in replicating cells. However, in these cells, a majority (50-90%) of P-TEFb is sequestered in the 7SK small nuclear (sn) RNA-nucleoprotein complex (RNP) as an enzymatically inactive form [12–14]. Cellular and environmental stimuli release P-TEFb from the 7SKsnRNP and activate CDK9, leading to the stimulation of transcriptional elongation [15]. It was long thought that this “P-TEFb-equilibrium” is a primary mechanism by which P-TEFb function is regulated [11, 16]. However, P-TEFb is also regulated by CycT1 protein levels, which are rapidly reduced in the absence of CDK9 [17]. In quiescent T cells and macrophages, which represent the bulk of the HIV latent reservoirs, levels of CycT1 are vanishingly low despite of abundant CycT1 transcripts [17, 18]. In all these cells, CycT1 is rapidly degraded post-translationally by cellular proteasome-dependent mechanism(s) [19].

Similar CycT1 degradation occurs in various in vitro models of unresponsive T cells, suggesting that low protein levels of functional P-TEFb play critical roles in impaired T cell responses in these cells [19]. We recently demonstrated that phosphorylation of CycT1 by the protein kinase C (PKC) promotes P-TEFb assembly in activated/proliferating cells, preventing CycT1 degradation. In resting T cells, CycT1 is dephosphorylated by the protein phosphatase 1 (PP1) and dissociated from CDK9, which causes rapid proteolysis of CycT1 [19]. We identified two threonine residues (T143 and T149) in CycT1 as the phosphorylation sites required for the tight interaction between CycT1 and CDK9 [19]. Moreover, we identified Seven in Absentia Homolog (Siah) 1/2 as E3 ligases that are responsible for the degradation of unphosphorylated and CDK9-unbound CycT1 [20].

Since P-TEFb is an essential cellular co-factor for HIV transcription, we reasoned that the absence of P-TEFb in quiescent CD4+ T cells is the main mechanism by which HIV latency is maintained in these cells. Thus, increasing P-TEFb protein levels in these cells is a mandatory step to control and prevent HIV latency. Based on our previous study, we designed mutant CycT1 proteins that are resistant to protein degradation and remain tightly bound to CDK9. One of the mutant proteins, named constitutively-active (CA)-CycT1 was fully functional and stably expressed in resting CD4+ T cells.

Establishment of HIV latency was significantly delayed in CD4+ T cells expressing CA-CycT1. These results present significant therapeutic potential for the development of effective therapies for HIV/AIDS and HIV-related malignancies.

## Results

### Designing mutant CycT1 proteins that interact stably with CDK9

We have previously demonstrated that two highly conserved threonine residues (T143 and T149) in the cyclin box region of CycT1 are phosphorylated by isoforms of PKC, which promotes the P-TEFb assembly and prevents CycT1 degradation (**Fig.1A**) [19]. Based on a three-dimensional structure of CycT1 [21], we conducted a computational simulation of the CDK9:CycT1 interaction surface with or without phosphorylation at T143 and T149 (**Fig.1B**). Phospho-T149 interacts with a basic residue in the N-terminus of CDK9 (R69), indicating that this threonine is involved in the intermolecular interaction between CycT1 and CDK9 [19]. Phospho-T143, on the other hand, faces towards the inside of the CycT1’s cyclin box structure and interacts with N-terminal residues in CycT1, indicating that it is involved in the intra-molecular interaction that stabilizes the cyclin box structure. Therefore, we first substituted these residues with glutamates, and examined whether the resulting mutant CycT1 (T143E/T149E) interacts with CDK9 by co-immunoprecipitation (**Fig.1A**). As shown previously, the phopho-negative mutant

**Figure 1.**
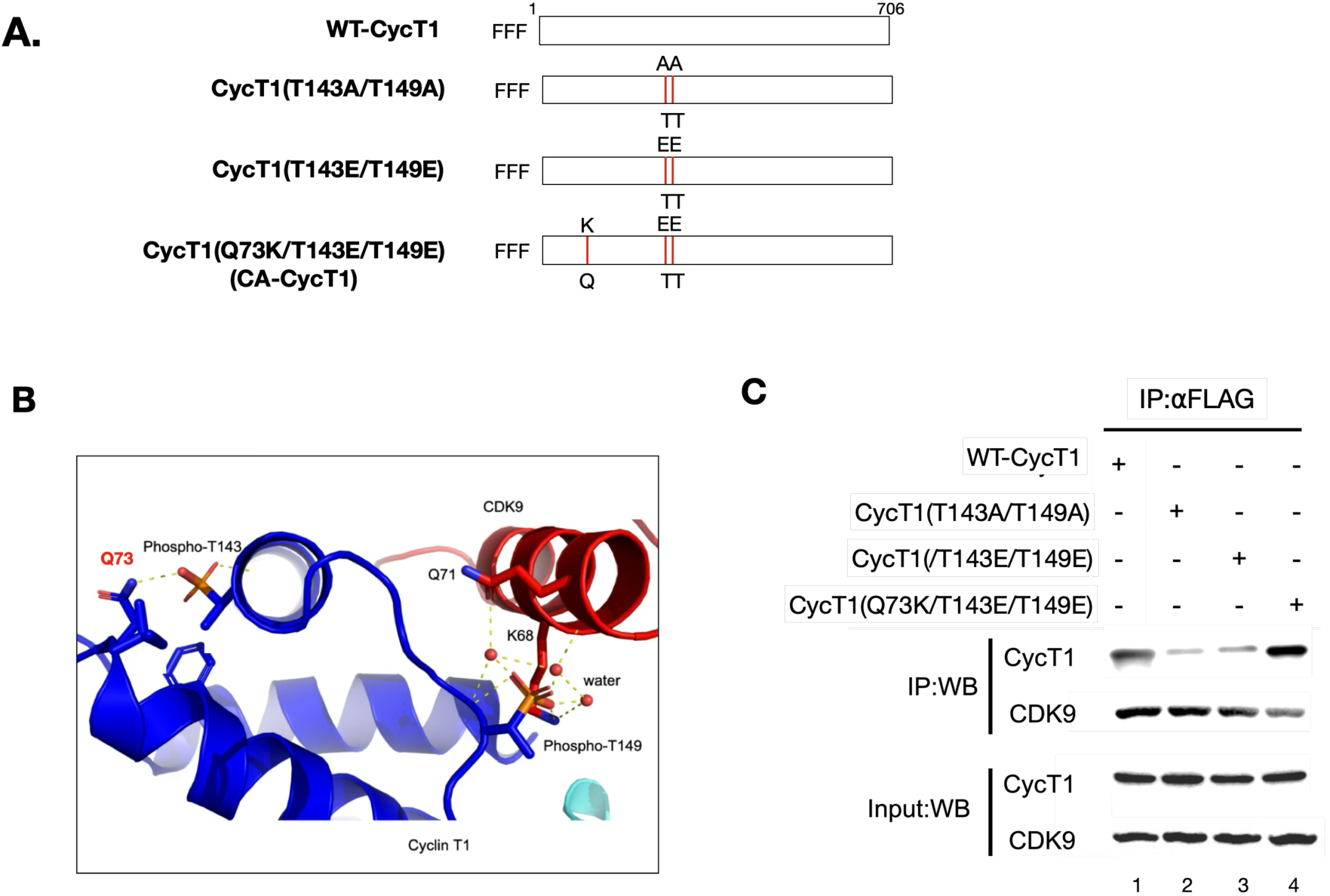
**A.** A schematic presentation of the WT and CA-CycT1 used in this study. The positions of point mutations are indicated by red lines. **B.** Model of P-TEFb where CycT1 is phosphorylated at T143 and T149. The model was created by adding phosphates to T143 and T149 in the published crystal structure of P-TEFb (PDB ID 3MI9), followed by energy minimization and molecular dynamics (MD) simulations. Residues predicted to interact with T143 and T149 are Q73 in CycT1 and K68 in CDK9, respectively. **C.** HA(h) epitope-tagged WT-CycT1 or indicated mutant CycT1 proteins and Flag (f) epitope-tagged CDK9 proteins were co-expressed in the presence of bortezomib (+/- signs on top) in 293T cells. CDK9 proteins were immunoprecipitated with anti-Flag antibody. CycT1 proteins co-immunoprecipitated with CDK9 were detected by anti-HA antibody (upper panels). Lower panels contain input levels of CycT1 and CDK9.

CycT1 (T143A/T149A) failed to interact with CDK9 (**Fig.1C**, lane 2). Although simply substituting these threonines with negatively-charged amino acids was expected to mimic the phosphorylation-dependent interaction of the wild type (WT) CycT1, these mutant CycT1 (T143E/T149E) proteins only partially reconstituted this interaction (**Fig.1C**, lane 3). The 3D model indicates that the side chain of the glutamine at the position 73 (Q73) is the closest to T143, suggesting that Q73 is an acceptor of the phospho-T143 (**Fig.1B**). Interestingly, adding a positive charge at this position by substituting the glutamine with lysine (Q73K) in CycT1 (T143E/T149E) almost fully restored the interaction between CDK9 and CycT1 (**Fig.1C**, lane 4), indicating that addition of a positive charge compensates for insufficient negative charges on E143 (E143 has one negative charge whereas naturally occurring phospho-T143 has two), tightening the intramolecular interaction between these two residues. We conclude that the mutant CycT1 (Q73K/T143E/T149E) is a phospho-mimetic mutant that can tightly interact with CDK9 (hence, named constitutively-active (CA)-CycT1).

### CA-CycT1 is functional

To confirm that the CA-CycT1 is functional, triple-Flag-epitope tagged (3xF) wild-type (WT) CycT1 and 3xF:CA-CycT1 proteins were expressed in the previously established CycT1-knockout (KO) HEK293 cells [22], in the presence of the reporter LTR luciferase (HIVLTR-Luc) gene with or without Tat [23]. Since Tat requires CycT1 to recruit P-TEFb to the HIV-LTR, Tat did not activate HIV transcription in CycT1-KO cells (**Fig.2A**, black bar 2). Transient expression of WT-CycT1 or CA-CycT1 in CycT1-KO cells restored Tat-dependent luciferase expression from the LTR to similar levels, which is equivalent to that achieved by endogenous P-TEFb in HEK293T cells (**Fig.2A**, black bar 3, compared with black bar 1). Western blotting confirmed that ectopically introduced WT-CycT1 and CA-CycT1 were expressed in CycT1-KO cells at levels similar to that of the endogenous CycT1 proteins in HEK293T cells (**Fig.2A**, bottom panels).

**Figure 2.**
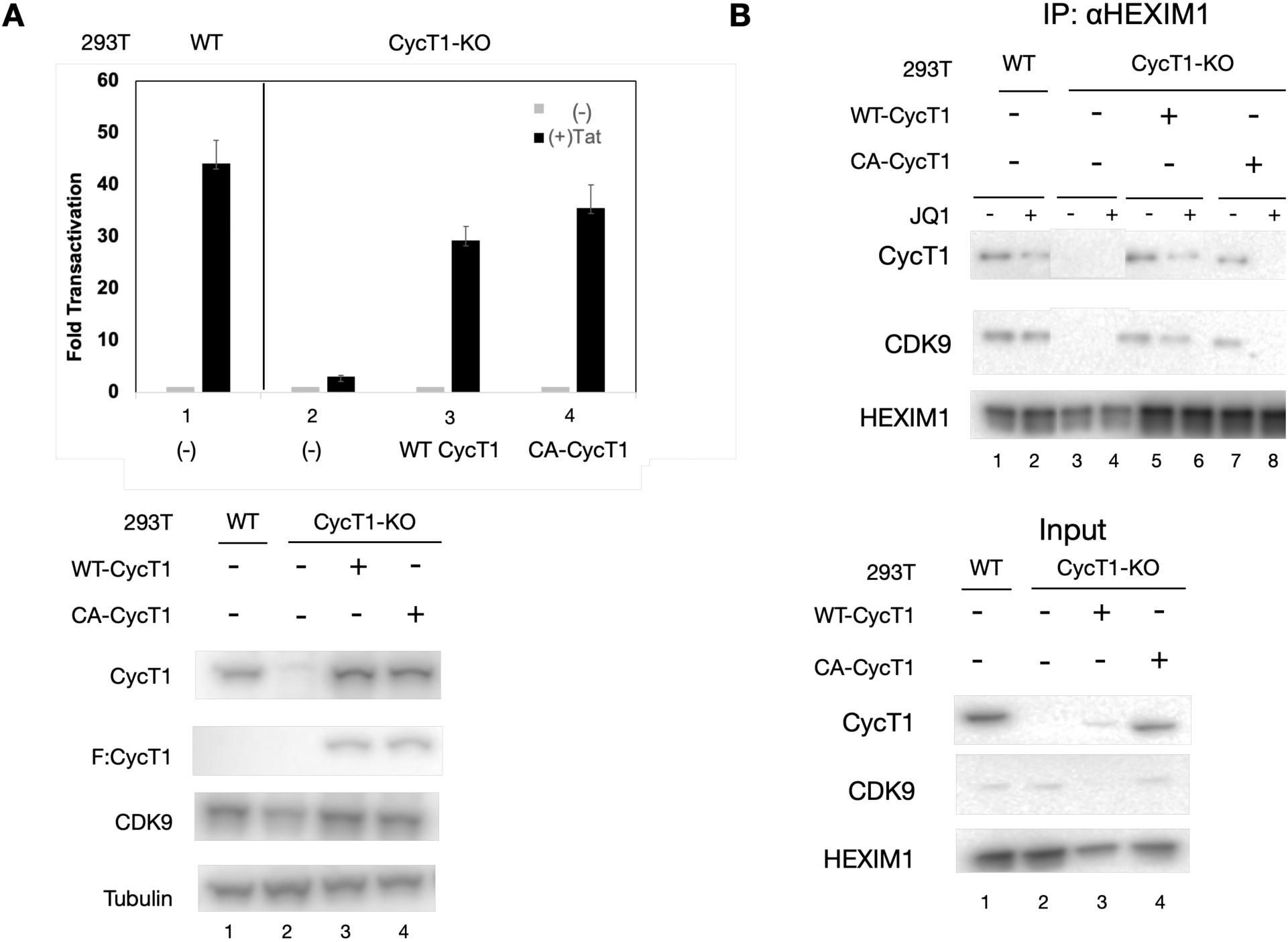
**A**. 293T or 293T/CycT1-KO cells were transfected with the reporter HIV-LTR-Luc plasmid with or without the plasmids encoding Tat or CycT1 (WT or CA). 48 hours after transfection, cells were harvested, and luciferase activities were measured. Data are presented as fold transactivation over values obtained without Tat. **B.** 293T or 293T CycT1-KO cells stably expressing WT or CA-CycT1 were treated with or without JQ1 for 1hr. Cells were lysed and CycT1 proteins were immunoprecipitated, followed by WB to detect HEXIM1 and CDK9. After extensive washes, proteins in immunoprecipitated were separated by SDS-PAGE and detected by western blotting with indicated antibodies, Input (∼10%) protein levels are also analyzed by western blotting (lower panels).

Next, we examined whether P-TEFb containing CA-CycT1 is incorporated into the 7SKsnRNP in unstimulated cells and released upon stimulation known to activate P-TEFb. 3xF:WT-CycT1 and 3xF:CA-CycT1 proteins were expressed in CycT1-KO cells. Cells were untreated or incubated with a BET-Bromodomain inhibitor (BETi), JQ1, for 1hr. The interaction between a 7SKsnRNP component HEXIM1 and P-TEFb was analyzed by co-immunoprecipitation. As shown in **Fig.2B**, in the control 293T cells, the endogenous CycT1 and CDK9 proteins were co-immunoprecipitated with HEXIM1 in the absence of JQ1 (**Fig.2B**, lane 1). The amount of CycT1 and CDK9 proteins associated with HEXIM1 were decreased upon JQ1 stimulation (**Fig.2B**, lane 2), confirming that JQ1 released P-TEFb from 7SKsnRNP as previously demonstrated [24]. Similarly, WT and CA-CycT1 expressed in CycT1-KO cells were associated with HEXIM1 (**Fig.2B**, lanes 3 and 5) in the absence of JQ1, and the levels of HEXIM1-associated WT and CA-CycT1 proteins were decreased upon JQ1 treatment (**Fig.2B**, lanes 4 and 6). These results indicate that ectopically expressed WT or CA-CycT1 function as endogenous CycT1 in the context of the association with and dissociation from the 7SKsnRNP. Taken all together, CA-CycT1 forms a stable and functional P-TEFb complex in cells.

#### CA-CycT1 is stably expressed in resting CD4+ T cells

Since our previous studies indicated that CycT1 tightly associated with CDK9 is resistant to protein degradation, we examined whether CA-CycT1 can be expressed stably in resting CD4+ T cells [19].

First, we established a rapid assay to evaluate mutant CycT1 proteins’ resistance to protein degradation in HEK293T cells (**Fig.S1**). WT and mutant CycT1 proteins were expressed in CycT1-KO cells. Cells were then treated with staurosporin at 3μM for 18 hours, which causes dephosphorylation and subsequent proteolysis of WT-CycT1, measured by western blotting. WT-CycT1 proteins were rescued by proteasomal inhibition by bortezomib at 10nM for 4hrs (**Fig.S1,** lanes 1 to 3). Levels of CA-CycT1 proteins, on the other hand, were not decreased by staurosporin (**Fig.S1,** lanes 4 and 5), indicating that CA-CycT1 is resistant to dephosphorylation-dependent proteolysis. Bortezomib did not increase levels of CA-CycT1 (**Fig.S1,** lanes 6).

Next, we examined whether CA-CycT1 proteins can be expressed stably in resting CD4+ T cells. VSV-G-pseudotyped lentiviruses encoding 3xF:WT-CycT1, CA-CycT1 or no open reading frame (“Empty”) were produced in HEK293 T cells, and infected into CD3/CD28-activated freshly isolated CD4+ T cells. Cells were put into a resting state by gradually reducing the growth factors (schematically described in **Fig.3A**). On indicated days post infection, small aliquots of cells were lysed, and levels of endogenous CycT1 proteins and ectopically expressed WT and CA-CycT1 proteins were monitored by western blotting. The anti-CycT1 (**Fig.3B**, top panels) antibodies detected the endogenous and ectopically-expressed CycT1 whereas the anti-Flag antibodies (**Fig.3B**, second panels) detected ectopically-expressed CycT1 only. As shown in **Fig. 3B**, levels of endogenous CycT1 proteins declined over time after removing growth factors (**Fig.3B**, top panels, lanes 1 to 4). Levels of ectopically introduced WT-CycT1 also declined at a similar rate to that of endogenous CycT1 (**Fig.3B**, second panel, lanes 5 to 8). In contrast, levels of ectopically expressed CA-CycT1 remained high (**Fig.3B**, second panel, lanes 9-12). Stable expression of ectopically expressed CA-CycT1 in resting CD4+ T cells was also confirmed by FACS using FITC-labelled anti-Flag (**Fig.3C**, upper panels) and FITC-labelled anti-CycT1 antibodies after cell permeabilization (**Fig.3C**, lower panel) [25]. It is to be noted that more than 80% of CD4+ T cells were transduced with lentiviruses encoding CA-CycT1, judged by Flag-positive cell population. Expression of an activated T cell-specific marker (HLA-DR) did not increase by the ectopic expression of WT or CA-CycT1 (**Fig.4**). These results indicate that CA-CycT1 forms a degradation-resistant P-TEFb expressed stably in resting CD4+ T cells.

**Figure 3.**
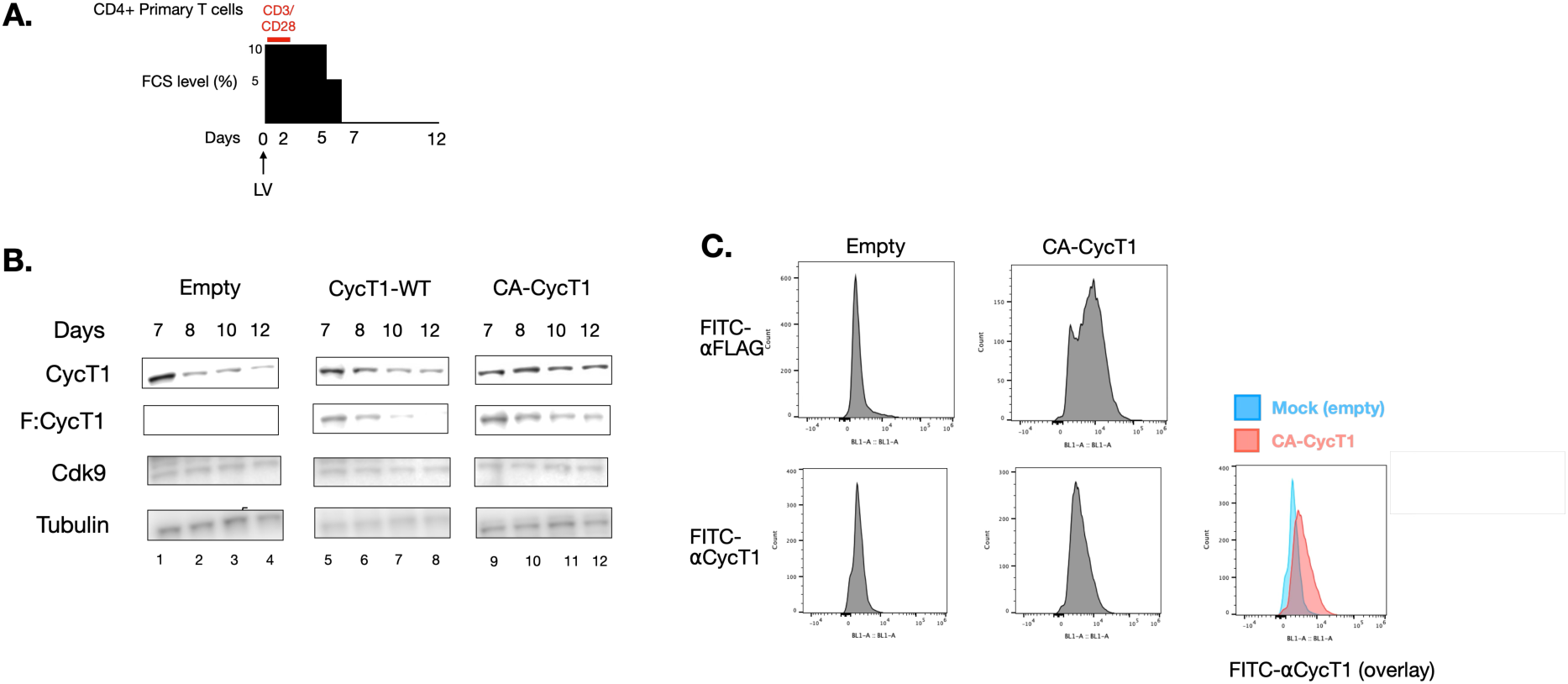
**A.** A diagram presents the experimental scheme. Primary T cells were stimulated with αCD3/αCD28 (red bar) on Day 0, followed by infection of VSV-G pseudotyped lentivirus (LV). Cells were gradually put into the resting states by reducing FCS and IL2. **B.** On indicated days after LV transduction, small aliquots of cells were spun down and lysed with SDS-sample buffer for western blotting with indicated antibodies. Tubulin was used as a loading control. **C.** The expression of endogenous and ectopically-expressed CycT1 proteins was also analyzed by flow cytometry. On indicated days after LV infection, small aliquots of cells were permeabilized and fixed, followed by incubation with FITC-labelled anti-CycT1 or anti-Flag antibodies.

### CA-CycT1 delays the establishment of HIV latency

Since CycT1 is an essential cellular co-factor for HIV Tat, lack of CycT1 in resting CD4+ T cells would represent a major contributor for maintaining the low HIV transcription levels in latently infected cells [7, 26]. Therefore, we examined whether ectopic expression of CA-CycT1 in these cells affects HIV latency. CD3/CD28-activated freshly isolated CD4+ T cells were infected with HIV Env-pseudotyped RGH dual-fluorescent-reporter viruses [27]. 24 hours after Env-RGH infection, samples were divided into two portions and infected with VSV-G-pseudotyped lentiviruses encoding CA-CycT1 or empty lentiviruses. Lentiviral-transduced cells were further enriched by puromycin treatment for 48 hours (**Fig. 5A**). HIV-infection was measured by the internal-EF-promotor-driven mCherry expression whereas HIV gene expression was measured by the LTR-driven eGFP expression. As shown in **Fig.5B**, higher rates of HIV-expressing cells were detected in CD4+T cells transduced with CA-CycT1 on day 13 post infection, compared to the control cells (**Fig.5B**, right panels). The time course analysis of eGFP positive cells confirmed that the establishment of HIV latency was significantly delayed in the cells expressing CA-CycT1 **(Fig.5C**). The ratios of HIV-positive cells between control and CA-CycT1 expressing cells indicate that 5x more cells are expressing HIV by CA-CycT1 over the course of infection (**Fig.5D**). These results indicate that the stable expression of CA-CycT1 in resting CD4+ T cells delay the establishment of HIV latency.

**Figure 5.**
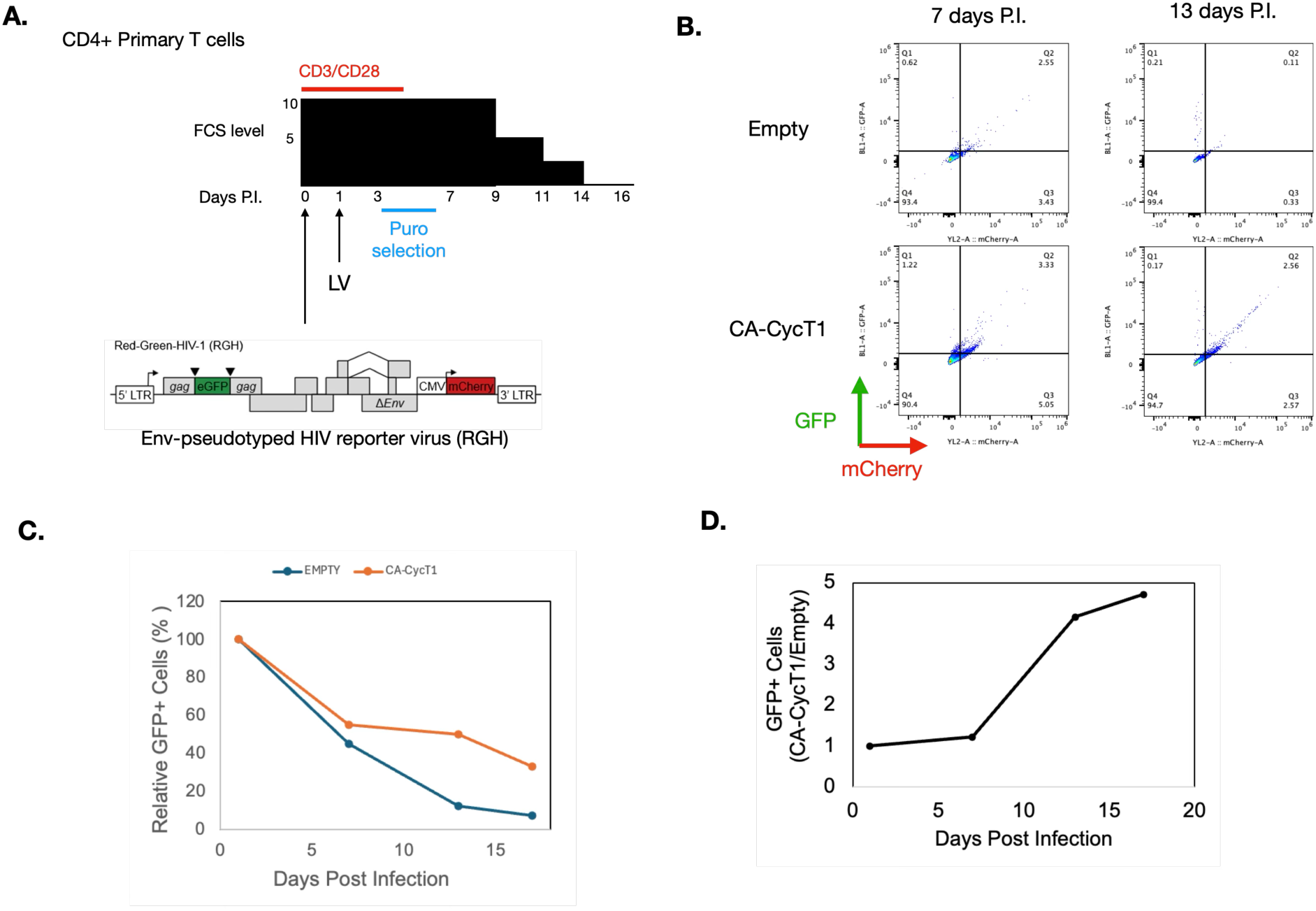
**A.** A diagram presents the experimental scheme. Primary T cells stimulated with αCD3/αCD28 were infected with HIV Env-pseudotyped RGH HIV dual-color reporter virus. 24 hours post infection, cells were divided into two portions and infected with VSV-G-pseudotyped lentivirus encoding CA-CycT1 or no proteins (LV). Lentivirally-transduced cells were enriched by puromycin treatment (5ng/mL) for 48 hrs. Cells were then slowly put into the resting state by gradually reducing stimulants. **B.** HIV-expressing (GFP+) and HIV-infected cells (mCherry+) were measured on the indicated days post infection by FACS analysis. **C.** Percentage of productively HIV-infected cells (GFP+/mCherry+) were plotted over days after infection. **D.** The ratios of HIV-positive cells between control and CA-CycT1-expressing cells were calculated on each day of measurement and plotted.

## Discussion

While the activity of P-TEFb is regulated primarily by its equilibrium with 7SKsnRNP in actively replicating cells, levels of functional P-TEFb are diminished in resting T cells because of CycT1 degradation. In this study, we designed mutant CycT1 proteins (called CA-CycT1) that are highly resistant to protein degradation. CA-CycT1 is fully functional as a P-TEFb component. When CA-CycT1 is expressed in resting CD4+ T cells, it is stably expressed even after levels of endogenous CycT1 proteins are diminished. Importantly, CA-CycT1 delays the establishment of HIV latency in CD4+ T cells without eliciting T cell activation. These results indicate that HIV latency can be controlled by manipulating cellular P-TEFb expression and function in resting CD4+ T cells.

Although CA-CycT1 maintains the elongation of HIV transcription at high levels, HIV latency could still occur. As previously demonstrated, latent HIV can be established in activated CD4+ T cells where P-TEFb is abundantly expressed [27, 28]. Various genetic and epigenetic mechanisms are involved in the establishment of HIV latency in these cells [29, 30]. In particular, HIV transcription can be severely silenced by transcription interference (TI) when HIV is integrated in actively transcribed genes [31, 32]. HIV transcription is blocked because on-going RNAPII transcribing the genes upstream or downstream (in reverse direction) of the integrated HIV reads through the boundary between the cellular gene and HIV, inhibiting the recruitment of RNAPII to TSS of HIV. Activation of HIV transcription requires a potent induction of RNAPII/P-TEFb recruitment to HIV TSS and inhibition of cellular transcription to occur simultaneously [31].

Therefore, it is necessary to specifically recruit P-TEFb to HIV loci without globally activating transcription elongation to reverse TI-mediated HIV latency [31]. In this scenario, increasing functional P-TEFb would be highly beneficial for activating TI-suppressed HIV. Since the main function of many currently used LRAs, including histone deacetylase inhibitors (HDACs) such as vorinostat or BET-bromodomain inhibitors (BETi), is to release P-TEFb from 7SKsnRNPs and activate it, it will be of interest to examine whether CA-CycT1 increases LRAs’ effects on reactivating latent HIV in these cells [33].

To our knowledge, this is the first example of expressing a high level of functional P-TEFb in resting CD4+ T cells, which modulates HIV transcription. Although P-TEFb is a major cellular transcription elongation factor required for many cellular genes, expressing high levels of CA-CycT1 in resting T cells had a minor effect on T cell physiology. Our data indicate that expression of T cell activation markers is unaffected by CA-CycT1 expression. Also, global gene expression analysis indicate only a small portion of genes are activated by CA-CycT1. Since a majority of P-TEFb is incorporated into the 7SKsnRNP and inactivated in even highly replicating cells, only a small amount of P-TEFb with CA-CycT1 will be enzymatically active and stimulate transcriptional elongation [13, 14]. Moreover, one of the immediate early genes responding to the active/free P-TEFb is its own inhibitor, HEXIM1, causing a rapid re-assembly of 7SKsnRNP and attenuating global transcription elongation [34, 35]. Since Tat can extract P-TEFb from 7SKsnRNP and stimulates HIV transcription, it is expected that increasing the amount of P-TEFb proteins in resting CD4+ T cells will be highly advantageous for specifically activating HIV transcription with minimum effects on cellular genes [36].

Even though this is a proof-of-concept study that increasing functional P-TEFb in resting CD4+T cells can affect HIV latency, questions remain as to how CA-CycT1 can be introduced in latently infected cells in clinical settings. While using lentiviral delivery system might not be very practical as a therapeutic agent, other methods including mRNA/nanoparticles are increasingly becoming a promising drug-delivery system. Another approach to increase functional P-TEFb in resting cells is to perturbate the cellular pathways that regulate CycT1 degradation. Particularly, PKC families play a critical role in the stability of CycT1 by phosphorylating it [19]. We have previously identified that T143 and T149 of CycT1 are phosphorylated by PKC [19]. Previous studies indicated that several isoforms of PKC are inactive or absent in resting cells [37–40]. In addition, HIV Tat no longer works when T cells are deprived of PKC by constant stimulation with PMA, which also promotes the proteolysis of CycT1 [41]. The activity of HIV Tat is restored when a constitutively active form of PKC is expressed in these cells [41]. Therefore, it is of interest to identify which isoform(s) of PKC are downregulated in HIV latently infected cells and test whether reinstalling the kinase activity is sufficient to increase CycT1 proteins and activate HIV transcription in these cells.

There are additional conditions, where low levels of P-TEFb play important roles, such as in resting or differentiated cells, anergic and exhausted T cells, or memory B/T cells [42]. We have demonstrated that CycT1 degradation occurs in various models of chronically stimulated T cells [19]. Lack of functional P-TEFb in these cells could result in the deficiency of producing proper cytokines responding to external stimuli. In addition, since the replication of many other viruses, especially integrated proviruses, also requires P-TEFb, the loss of P-TEFb also contributes to their latent infection and viral persistence by escaping from immune surveillance [43]. The presented data will inform how they could be manipulated for optimal responses, be it in wound healing, inflammation, immune responses or improved general function.

The regulatory mechanism of this CycT1 protein turnover represents broad implications for a variety of fields ranging from HIV biology to cancer immunology. Our data represent significant potential to establish systems for controlling disease onsets by fine-tuning cellular transcriptional networks via manipulating the protein levels of critical cellular transcription factors. Finally, since P-TEFb is required for transcription mediated by at least two (c-Myc and Sox2) of the “Yamanaka factors”, we envision that even reprogramming, as occurs with iPS cells, could be improved by informed manipulation of P-TEFb homeostasis [44–47].

## Materials and Methods

### Cell Culture

Human Embryonic Kidney (HEK) 293T cells were cultured in Dulbecco’s modified Eagle’s medium (DMEM) (Corning) with 10% fetal bovine serum (FBS) (Life Technologies). Human primary blood mononuclear cells (PBMC) were isolated from LSR cones (STEMCELL Technologies) by Lymphoprep reagent (STEMCELL Technologies) and maintained in Roswell Park Memorial Institute (RPMI)1640 (Corning) supplemented with 10%FCS, L-Glutamate, and sodium pyruvate (cRPMI). CD4+ T cells were further isolated by negative selection using EasySep Human CD4+T Cell Isolation Kit (STEMCELL Technologies) and maintained in cRPMI. Cells were expanded by CD3/CD28 stimulation (ImmunoCult Human CD3/CD28 T Cell activator, STEMCELL Technologies) for up to one week for downstream experiments. After stimulations cells were maintained in cRPMI with recombinant IL2 (Sigma).

### Antibodies

Antibodies used in this study are listed below.

| Target | label | Species | Clone# | Cat# |
| --- | --- | --- | --- | --- |
| CycT1 |  | Mouse | E3 | SCBT/sc-271348 |
| CycT1 | FITC | Mouse | E3 | SCBT/sc-271348 FITC |
| Flag |  | Mouse | M2 | SIGMA/F-3165 |
| Flag | FITC | Mouse | M2 | SIGMA/F-4049 |
| Tubulin |  | Rabbit |  | Proteintech/11224-1-AP |
| HEXIM1 |  | Rabbit |  | Proteintech/15676-1-AP |
| CDK9 |  | Rabbit |  | Proteintech/11705-1-AP |
| CD25 | FITC | Mouse |  | BD/ 555431 |
| CD69 | PE | Mouse |  | BD/555531 |
| normal IgG | FITC | Mouse |  | SCBT/sc-516614 |
| normal IgG | PE | Mouse |  | SCBT/sc-2857 |

### Lentiviral Production

High-titre stock solutions of VSV-G-pseudotyped lentiviruses were produced by transfection in early passage (P<20) 293T cells, following the protocol established in the Marson lab [48, 49]. Briefly, lentiviral vectors, pMD.2g, and psPAX2 were transfected into 293T cells by Lipofectamine 3000 (Life Technologies). 6 to 8hrs after transfection, media were replaced with OptiMEM supplement with GlutaMAX Supplement (Invitrogen), 5% FCS, 1 mM sodium pyruvate (Fisher Scientific), and 1×MEM nonessential amino acids (Fisher Scientific) (cOPTI) and 1x Viral Boost (Alstem Bio). 24 and 48hrs after transfection, culture supernatants were transferred, and cell debris were removed by brief centrifugation (500xG, 5min). Lentiviruses in the supernatant were precipitated by Lenti-X Concentrator (Takara), resuspended with ∼1mL cRPMI, aliquoted, and stored in at -75 ℃ HIV-Env-pseudotyped HIV reporter viruses were produced similarly, by co-transfecting HIV Env expression plasmids, instead of pMD.2g.

### Western Blotting

Approx.10^6 cells were pelleted and lysed directly with 2xLaemmli SDS sample buffer (BioRad) and heated at 95C for 10min. Approx. one-fifth of the lysates were separated in 4-12% SDS-PAGE gels (Genscript) in Tris-MOPS-based running buffer, and transferred electronically to PVDF membranes. After blocking with 5% non-fat dry milk or 3% BSA, the membranes were incubated with indicated primary antibodies, followed by incubation with HRP-conjugated secondary antibodies with extensive washing.

Positive bands were visualized by SuperSignal West Femto Maximum Sensitivity Substrate (ThermoFisher) and LiCor Image analyzer. When necessary, the signals were enhanced by Can-Get-Signal Immunoreaction Enhancer Solution (TOYOBO). Band intensities were quantified by LiCOR imaging analyzer.

### Flow Cytometric Analysis

Endogenous or ectopically expressed CycT1 proteins were also measured by flow cytometry (ATTUNE) [25]. Cells were fixed and permeabilized by using the Fox-P Staining Kit (eBioscience) by following the manufacturer’s instruction. Cells were incubated with FITC-labelled normal mouse IgG (negative control), FITC-labelled anti-CycT1 or FITC-labelled anti-Flag for 30 min.

## Statistical analysis

Experiments using primary T lymphocytes were performed with at least six different blood donors. Presented data are representative results from three independent experiment, all experiments were done in duplicate. Thus, bars represent SEM, n=3. A Student’s *t*-test was preformed to measure the significance of the data (* P<0.05, ** P<0.01, *** P<0.001).

## Acknowledgement

We thank Dr. Ujjwal Rathore for providing helpful instructions for producing high-titre lentiviruses and sharing protocols for LV infection to primary CD4+ T cells. We also thank Dr. Alan D. Frankel and the members of the Frankel lab for helpful discussions. We thank Drs. Yang Li and Yusuke Matsui for providing plasmids. We thank Zeping Luo, Dan Irwin, and Dr. Aung Chein for technical assistance. Dr. B. Matija Peterlin for extensive advices. Dr. Ants Fujinaga for editing and proofreading the manuscript. This study was supported by

---

**Figure S1.**
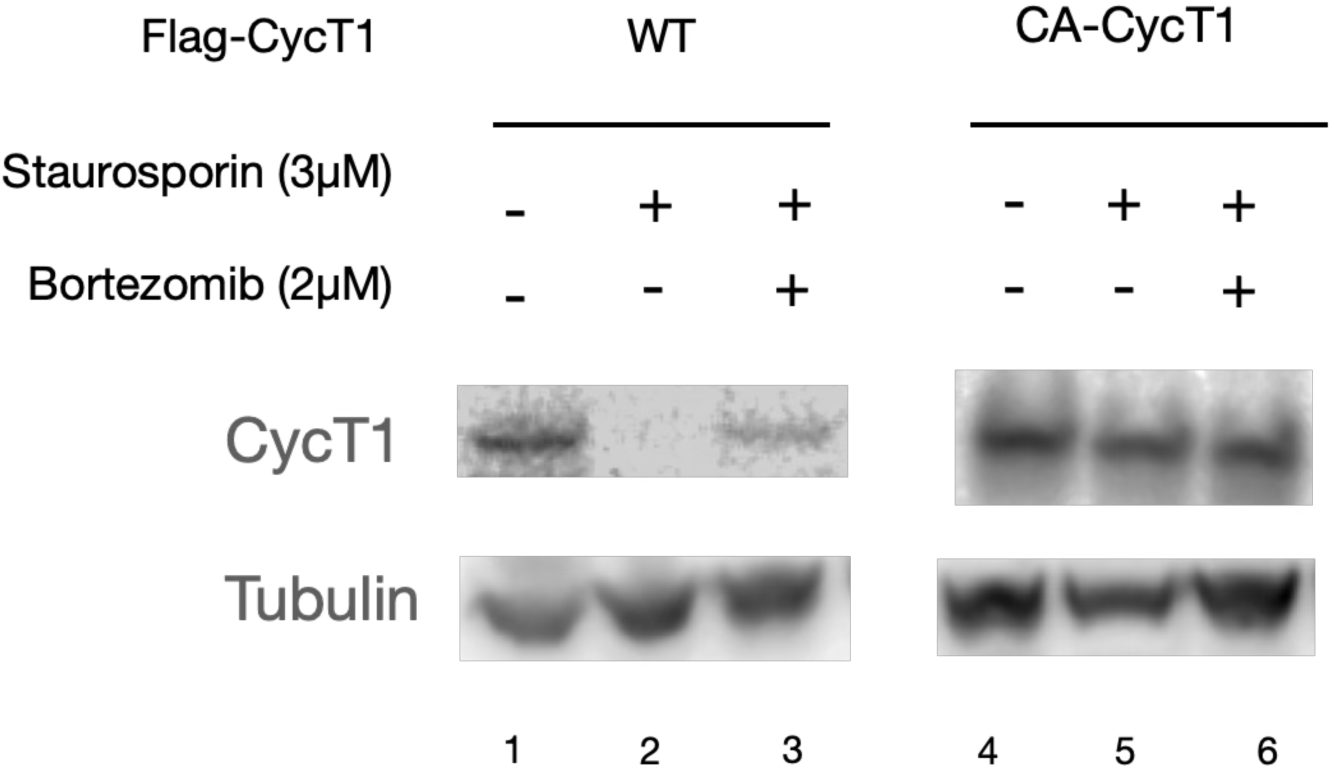
CycT1-KO cells stably expressing WT- or CA-CycT1 proteins were treated with indicated the concentrations of staurosporin for 18 hours with or without bortezomib for 4hrs. Cells were collected and lysed. CycT1 and tubulin proteins were detected by western blotting.

## References

1. Deeks, S.G., et al., International AIDS Society global scientific strategy: towards an HIV cure 2016. Nat Med, 2016. 22(8): p. 839–50.

2. Henrich, T.J., et al., HIV-1 persistence following extremely early initiation of antiretroviral therapy (ART) during acute HIV-1 infection: An observational study. PLoS Med, 2017. 14(11): p. e1002417.

3. Li, Z., et al., Reiterative Enrichment and Authentication of CRISPRi Targets (REACT) identifies the proteasome as a key contributor to HIV-1 latency. PLoS Pathog, 2019. 15(1): p. e1007498.

4. Kimata, J.T., A.P. Rice, and J. Wang, Challenges and strategies for the eradication of the HIV reservoir. Curr Opin Immunol, 2016. 42: p. 65–70.

5. Siliciano, J.D. and R.F. Siliciano, Recent developments in the effort to cure HIV infection: going beyond N = 1. J Clin Invest, 2016. 126(2): p. 409–14.

6. Cary, D.C., K. Fujinaga, and B.M. Peterlin, Molecular mechanisms of HIV latency. J Clin Invest, 2016. 126(2): p. 448–54.

7. Mbonye, U. and J. Karn, Transcriptional control of HIV latency: cellular signaling pathways, epigenetics, happenstance and the hope for a cure. Virology, 2014. 454–455: p. 328-39.

8. Ott, M., M. Geyer, and Q. Zhou, The control of HIV transcription: keeping RNA polymerase II on track. Cell Host Microbe, 2011. 10(5): p. 426–35.

9. Karn, J., Tackling Tat. J Mol Biol, 1999. 293(2): p. 235–54.

10. AJ, C.Q., A. Bugai, and M. Barboric, Cracking the control of RNA polymerase II elongation by 7SK snRNP and P-TEFb. Nucleic Acids Res, 2016. 44(16): p. 7527–39.

11. Fujinaga, K., F. Huang, and B.M. Peterlin, P-TEFb: The master regulator of transcription elongation. Mol Cell, 2023. 83(3): p. 393–403.

12. Peterlin, B.M., J.E. Brogie, and D.H. Price, 7SK snRNA: a noncoding RNA that plays a major role in regulating eukaryotic transcription. Wiley Interdiscip Rev RNA, 2012. 3(1): p. 92–103.

13. Zhou, Q. and J.H. Yik, The Yin and Yang of P-TEFb regulation: implications for human immunodeficiency virus gene expression and global control of cell growth and differentiation. Microbiol Mol Biol Rev, 2006. 70(3): p. 646–59.

14. Diribarne, G. and O. Bensaude, 7SK RNA, a non-coding RNA regulating P-TEFb, a general transcription factor. RNA Biol, 2009. 6(2): p. 122–8.

15. Fujinaga, K., P-TEFb as A Promising Therapeutic Target. Molecules, 2020. 25(4).

16. Egloff, S., CDK9 keeps RNA polymerase II on track. Cell Mol Life Sci, 2021. 78(14): p. 5543–5567.

17. Ramakrishnan, R., et al., Making a Short Story Long: Regulation of P-TEFb and HIV-1 Transcriptional Elongation in CD4+ T Lymphocytes and Macrophages. Biology (Basel), 2012. 1(1): p. 94–115.

18. Marshall, R.M., et al., Cyclin T1 expression is regulated by multiple signaling pathways and mechanisms during activation of human peripheral blood lymphocytes. J Immunol, 2005. 175(10): p. 6402–11.

19. Huang, F., et al., Reversible phosphorylation of cyclin T1 promotes assembly and stability of P-TEFb. Elife, 2021. 10.

20. Huang, F., et al., P-TEFb is degraded by Siah1/2 in quiescent cells. Nucleic Acids Res, 2022. 50(9): p. 5000–5013.

21. Tahirov, T.H., et al., Crystal structure of HIV-1 Tat complexed with human P-TEFb. Nature, 2010. 465(7299): p. 747-51.

22. Leoz, M., et al., HEXIM1-Tat chimera inhibits HIV-1 replication. PLoS Pathog, 2018. 14(11): p. e1007402.

23. Li, Y., et al., Enhanced NF-kappaB activation via HIV-1 Tat-TRAF6 cross-talk. Sci Adv, 2024. 10(3): p. eadi4162.

24. Bartholomeeusen, K., et al., Bromodomain and extra-terminal (BET) bromodomain inhibition activate transcription via transient release of positive transcription elongation factor b (P-TEFb) from 7SK small nuclear ribonucleoprotein. J Biol Chem, 2012. 287(43): p. 36609–16.

25. Couturier, J., et al., Regulation of cyclin T1 during HIV replication and latency establishment in human memory CD4 T cells. Virol J, 2019. 16(1): p. 22.

26. Mbonye, U., et al., Biogenesis of P-TEFb in CD4+ T cells to reverse HIV latency is mediated by protein kinase C (PKC)-independent signaling pathways. PLoS Pathog, 2021. 17(9): p. e1009581.

27. Dahabieh, M.S., et al., A doubly fluorescent HIV-1 reporter shows that the majority of integrated HIV-1 is latent shortly after infection. J Virol, 2013. 87(8): p. 4716–27.

28. Chavez, L., V. Calvanese, and E. Verdin, HIV Latency Is Established Directly and Early in Both Resting and Activated Primary CD4 T Cells. PLoS Pathog, 2015. 11(6): p. e1004955.

29. Churchill, M.J., et al., HIV reservoirs: what, where and how to target them. Nat Rev Microbiol, 2016. 14(1): p. 55–60.

30. Sengupta, S. and R.F. Siliciano, Targeting the Latent Reservoir for HIV-1. Immunity, 2018. 48(5): p. 872–895.

31. Lenasi, T., X. Contreras, and B.M. Peterlin, Transcriptional interference antagonizes proviral gene expression to promote HIV latency. Cell Host Microbe, 2008. 4(2): p. 123–33.

32. Han, Y., et al., Orientation-dependent regulation of integrated HIV-1 expression by host gene transcriptional readthrough. Cell Host Microbe, 2008. 4(2): p. 134–46.

33. Darcis, G., et al., An In-Depth Comparison of Latency-Reversing Agent Combinations in Various In Vitro and Ex Vivo HIV-1 Latency Models Identified Bryostatin-1+JQ1 and Ingenol-B+JQ1 to Potently Reactivate Viral Gene Expression. PLoS Pathog, 2015. 11(7): p. e1005063.

34. Liu, P., et al., Release of positive transcription elongation factor b (P-TEFb) from 7SK small nuclear ribonucleoprotein (snRNP) activates hexamethylene bisacetamide-inducible protein (HEXIM1) transcription. J Biol Chem, 2014. 289(14): p. 9918–25.

35. Tan, J.L., et al., Stress from Nucleotide Depletion Activates the Transcriptional Regulator HEXIM1 to Suppress Melanoma. Mol Cell, 2016. 62(1): p. 34–46.

36. Barboric, M., et al., Tat competes with HEXIM1 to increase the active pool of P-TEFb for HIV-1 transcription. Nucleic Acids Res, 2007. 35(6): p. 2003–12.

37. Gould, C.M. and A.C. Newton, The life and death of protein kinase C. Curr Drug Targets, 2008. 9(8): p. 614–25.

38. Li, Y., et al., Protein Kinase C Family: Structures, Biological Functions, Diseases, and Pharmaceutical Interventions. MedComm (2020), 2025. 6(11): p. e70474.

39. Mann, T.H., Knox, H.M. et al, Different signaling interpretations by PKC eta and theta control T cell functionand exhaustion. BioRxiv, 2024.

40. Way, K.J., E. Chou, and G.L. King, Identification of PKC-isoform-specific biological actions using pharmacological approaches. Trends Pharmacol Sci, 2000. 21(5): p. 181–7.

41. Jakobovits, A., A. Rosenthal, and D.J. Capon, Trans-activation of HIV-1 LTR-directed gene expression by tat requires protein kinase C. EMBO J, 1990. 9(4): p. 1165–70.

42. Heissmeyer, V., et al., Calcineurin imposes T cell unresponsiveness through targeted proteolysis of signaling proteins. Nat Immunol, 2004. 5(3): p. 255–65.

43. Zaborowska, J., N.F. Isa, and S. Murphy, P-TEFb goes viral. Bioessays, 2016. 38 Suppl 1: p. S75–85.

44. Takahashi, K. and S. Yamanaka, Induction of pluripotent stem cells from mouse embryonic and adult fibroblast cultures by defined factors. Cell, 2006. 126(4): p. 663–76.

45. Arter, J. and M. Wegner, Transcription factors Sox10 and Sox2 functionally interact with positive transcription elongation factor b in Schwann cells. J Neurochem, 2015. 132(4): p. 384–93.

46. Eberhardy, S.R. and P.J. Farnham, Myc recruits P-TEFb to mediate the final step in the transcriptional activation of the cad promoter. J Biol Chem, 2002. 277(42): p. 40156–62.

47. Kanazawa, S., et al., c-Myc recruits P-TEFb for transcription, cellular proliferation and apoptosis. Oncogene, 2003. 22(36): p. 5707–11.

48. Carnevale, J., et al., RASA2 ablation in T cells boosts antigen sensitivity and long-term function. Nature, 2022. 609(7925): p. 174-182.

49. Rathore, U., et al., Systematic discovery of pro- and anti-HIV host factors in primary human CD4+ T cells. Cell, 2026. 189(12): p. 3651–3668 e31.

